# Engineering sensory ganglion multicellular system to model tissue nerve ingrowth

**DOI:** 10.1101/2023.01.30.526224

**Authors:** Junxuan Ma, Riccardo Tognato, Janick Eglauf, Sibylle Grad, Mauro Alini, Tiziano Serra

## Abstract

Discogenic pain is associated with deep nerve ingrowth in annulus fibrosus tissue (AF) of intervertebral disc (IVD). To model AF nerve ingrowth, primary bovine dorsal root ganglion (DRG) micro-scale tissue units are spatially organized around an AF explant by mild hydrodynamic forces within a collagen matrix. This results in a densely packed multicellular system mimicking the native DRG tissue morphology and a controlled AF-neuron distance. Such a multicellular organization is essential to evolve populational-level cellular functions and *in vivo*-like morphologies. Pro-inflammatory cytokine-primed AF demonstrates its neurotrophic and neurotropic effects on nociceptor axons. Both effects depend on the AF-neuron distance underpinning the role of recapitulating inter-tissue/organ anatomical proximity when investigating their crosstalk. This is the first *in vitro* model studying AF nerve ingrowth by engineering mature and large animal tissues in a morphologically and physiologically relevant environment. Our new approach can be used to biofabricate multi-tissue/organ models for untangling pathophysiological conditions and develop novel therapies.

## 1. Introduction

Sensory nerve ingrowth has been observed in the recovery from injury and in many degenerative diseases. In the healing of bone^[1]^ and tendon,^[2]^ sensory nerve ingrowth into the regenerating tissue may actively participate in tissue repair. On the contrary, aberrant innervations in joints^[3]^, intervertebral disc (IVD)^[4]^, and non-healed bone^[5]^ are frequently associated with chronic musculoskeletal pain.

The IVD is a fibrocartilaginous spacer in the spine and is normally devoid of nerves.^[6]^ Nerve ingrowth into IVD has been associated with chronic low back pain.^[4]^ This raises the hypothesis that the neo-innervation enables the IVD to generate pain (discogenic pain). Indeed, the axons growing into the annulus fibrosus (AF) tissue of disrupted IVD are labelled with calcitonin gene related peptide (CGRP) and substance P (SP)^[7]^ which are markers of the damage-sensing and pain-initiating neurons (*i*.*e*., nociceptors).^[8]^ Cell bodies of nociceptors locate in the dorsal root ganglion (DRG), an oval tissue within the spinal intervertebral foramina proximal to the IVD.^[9]^

Despite the clear association between sensory nerve ingrowth and discogenic pain, little is known about the biological role and mechanism of the AF nerve ingrowth. This points to an unmet need of experimental models. Currently, only *in vivo* rodent models have been reported to characterize sensory nerve ingrowth in IVD.^[10]^ However, the experimental manipulation and visualization of the tiny and complex axonal structures *in vivo* is challenging. Additionally, many rodent models are poor in predicting human diseases. More than 85% of their predicted therapeutic agents failed in human clinical trials.^[11]^ Mice and human have a largely divergent gene expression profile and regulatory program, although they do share the same set of genes.^[12]^ It has been generally recommended that, besides rodent models, therapeutic agents should be tested in large animal models before clinical trials.^[13]^ Nevertheless, large animal models are expensive, have low throughputs, and raise ethical concerns. Recently, human induced pluripotent stem cell (iPSC)-derived organoids are developed to faithfully mimic functional human organs but are still limited in studying multi-organ communication (*e*.*g*., between nerve and target tissue).^[14]^ The stepwise differentiation protocol to develop human neural organoid takes months. Moreover, the complexity of the nervous system containing various cell types (different types of neurons, glial and vessel cells etc.) is not yet fully recapitulated.^[15]^

Currently, to the best of our knowledge, there is not *in vitro* model to study the AF sensory nerve ingrowth. The challenge is to culture target tissue (*i*.*e*., AF) and DRG in a physiologically and morphologically relevant condition. Firstly, DRG neurons surrounded by satellite glial cell (SGC) envelope^[9]^ are densely packed in a multicellular architecture.^[16]^ This close spatial relationship is essential for their physiological crosstalk.^[17]^ Secondly, the model needs to recapitulate the close anatomical proximity between DRG and target tissue. The distance between nerve and target tissue may influence their communication.

Modern bioprinting allows microscale-precise positioning of cells/tissues where cells are usually sparsely distributed in a biocompatible hydrogel-based ink ^[18]^. During bioprinting, cells are subjected to a variety of stimuli that can result in cell damage or death.^[19]^ Importantly, DRG neurons’ role is to sense environmental stimuli.^[20]^ Therefore, they may be ill-suited for the common bioprinting processes. Recently, acoustic bioassemblies emerge as rapid, contactless and mild strategies to biofabricate physiologically relevant tissues.^[21]^ These techniques coordinate the assembly of biological components through the emergence of specific fluid patterns (*i*.*e*., pressure fields, surface instabilities/waves) upon excitatory external stimuli. The precise control of the external stimuli (*i*.*e*., frequency and amplitude of chamber vibration) generate hydrodynamic forces that orchestrate the assembly of biological components in predetermined morphologies.^[21]^ Once assembled, biological components are in tight cell-to-cell contact, dominant requirement to translate from morphed tissues-like structures to functional tissues!^[14b]^

We use Faraday waves^[22]^ to assemble large animal DRG micro-scale tissue units into densely packed architecture to which we refer to as multicellular system (Figure1a). The resultant DRG multicellular system is positioned in the predetermined ring-shaped geometry surrounding the AF tissue explant in a collagen matrix. We demonstrate the functional intercellular crosstalk and the structural self-organization evolved in the cultured DRG multicellular system.

**Figure 1.**
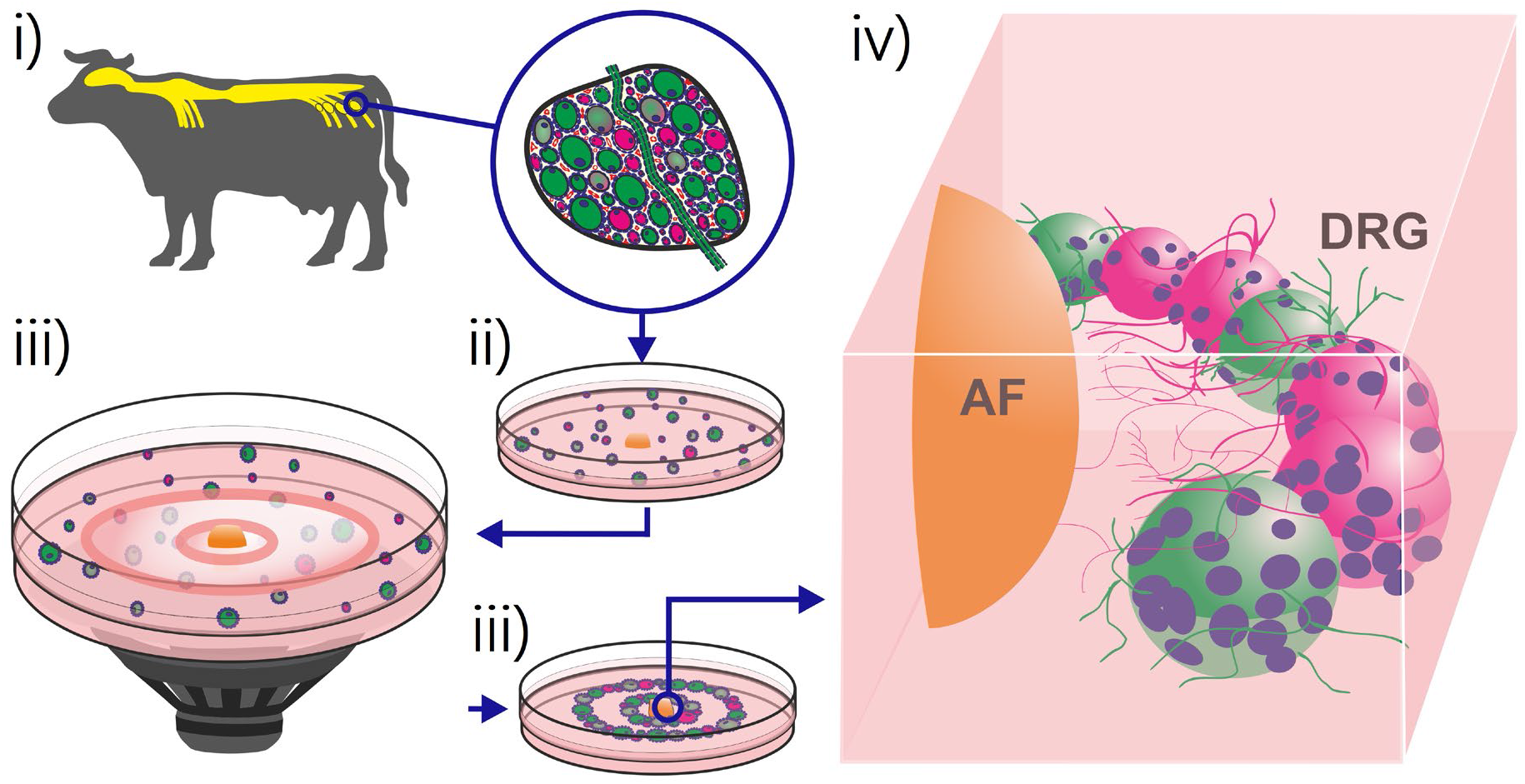
Schematic representation of the experimental approach used to engineer the sensory ganglion multicellular system. Bovine DRG (i) is disintegrated into micro-scale tissue units (ii) and assembled (iii) in ring-shaped multicellular systems surrounding the annulus fibrosus (AF) explant (iv).

The tissue nerve ingrowth is steered by two key mechanisms. The first is the neurotrophic (growth-promoting) effect which takes the role of elongating axons. Nevertheless, a simple elongation is not enough without a correct growing direction. Hence, the neurotropic (guidance) effect is of equal importance.^[23]^ We evaluated the influence of AF on these two mechanisms of axonal growth in the multicellular system as a proof of concept that pathophysiological nerve ingrowth can be engineered *in vitro*.

## 2. Results

### 2.1. Fabrication of DRG multicellular system surrounding the AF explant

The timeline of AF nerve ingrowth model fabrication is depicted in Figure 2a. The fabrication process starts with a pro-inflammatory cytokine priming of AF tissue explant to mimic the discogenic pain microenvironment. The applied cytokines are interleukin-1beta (IL-1β) and tumor necrosis factor alpha (TNF-α). These are well known to be associated with clinical discogenic pain.^[24]^ The cytokine-treated bovine AF explant cultured *ex vivo* shows high viability (Figure S1).

**Figure 2.**
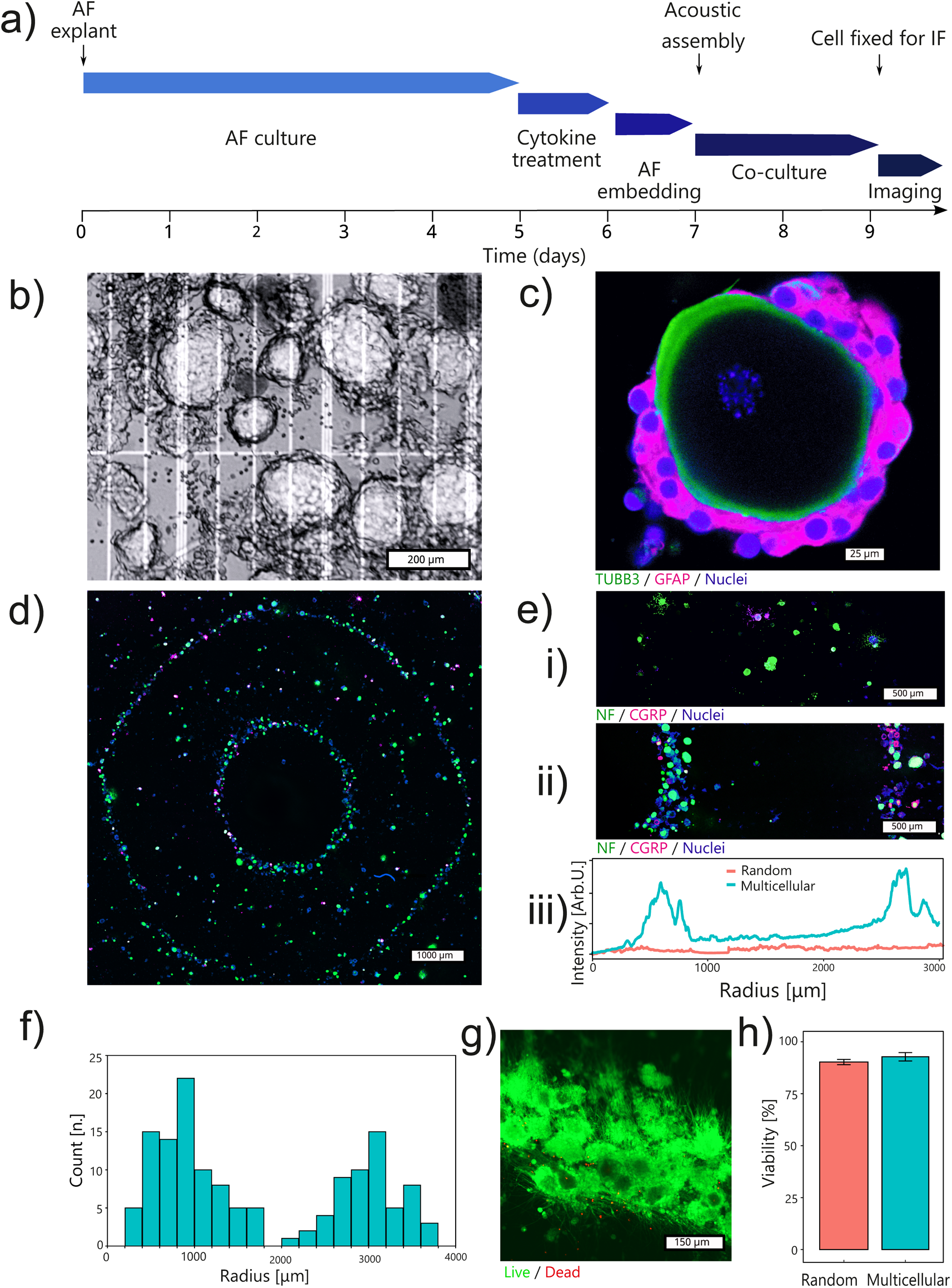
Establishment of the *in vitro* model to study the AF nerve ingrowth. **a)** Experimental timeline. **b)** Phase contrast image of enzymatically dissociated bovine DRG micro-scale tissue units (trypan blue staining). **c)** The mild enzymatic procedure preserves the envelope structure of native DRG tissue where neurons stained by tubulin beta 3 class III (TUBB3) are attached by satellite glial cells stained by glial fibrillary acidic protein (GFAP). **d-e)** Hydrodynamic forces assemble DRG micro-scale tissue units from a random spatial distribution (i) to a well-defined geometry (ii and iii). **f)** This assembly process leads to defined neurons-AF distances. **g-h)** The assembly procedure is mild. Viability of the assembled multicellular structure remained above 90% after 2 days of culture.

Next, we enzymatically disintegrate the whole DRG explant into micro-scale tissue units with high viability (87.1% by trypan blue staining, averaged from 24 independent experiments) (Figure 2b). Importantly, the products of enzymatical disintegration are not single cells but tissue units. The satellite glial cells (SGCs) ‘envelopes’ surrounding the neurons are preserved in the tissue units (Figure 2c). This structure is known to underly the neuron-glia communication.^[17b]^

To fabricate the AF-DRG system, the AF explant is placed in the center of a culture frame with the DRG tissue units dispensed around. This is followed by assembling the DRG tissue units into ring-shape geometry using hydrodynamic drag forces generated by a vertical vibration (Figure 2d). The ring diameters which determine the AF-DRG distance (Figure 2e (ii and iii)) are consistent among different independent experiments (Figure S2). Seeding DRG tissue units without applying hydrodynamic forces leads to their random positioning (Figure 2e). Neurons in the smallest ring are defined as ‘AF-close’ neurons. The ‘AF-close’ neurons are enriched at 0.8 ± 0.4 mm from the AF boarder. Neurons in the second ring serve as an ‘AF-far’ control and their distance to the AF boarder is 3.2 ± 0.8 mm (Figure 2f). The mild assembling process does preserve cell viability. Live / dead staining after 2 days of culture exhibits 92.8% viability in the assembled multicellular system which is at the same level as the random culture (90.3%) (Figure 2 g-h).

### 2.2. Functional characterization of the DRG multicellular system

Potassium-induced depolarization is commonly used to evaluate the functional maturation of neurons in a developing neural organoid.^[25]^ Rising the extracellular potassium chloride concentration to 50 mM causes a 48-mV membrane depolarization in neurons. This leads to the calcium influx through L-type voltage gated calcium channels (VSCCs) that can be detected through calcium imaging.^[25b]^ The maturity of neurons can be evaluated by this potassium depolarization-caused calcium influx. The functionality of several neural organoids has been evaluated following this method suggesting that 18-30 days of differentiation are needed for neurons to become functionally mature.^[25a, 26]^ In our model, the DRG tissues are obtained from adult cattle and the DRG neurons are readily functionally mature. Indeed, fast calcium upsurge in neurons following potassium depolarization is observed in both random culture and the multicellular system at day 2 (Figure 3a-b). This proves that they are functionally mature without the need of long-term differentiation.

**Figure 3.**
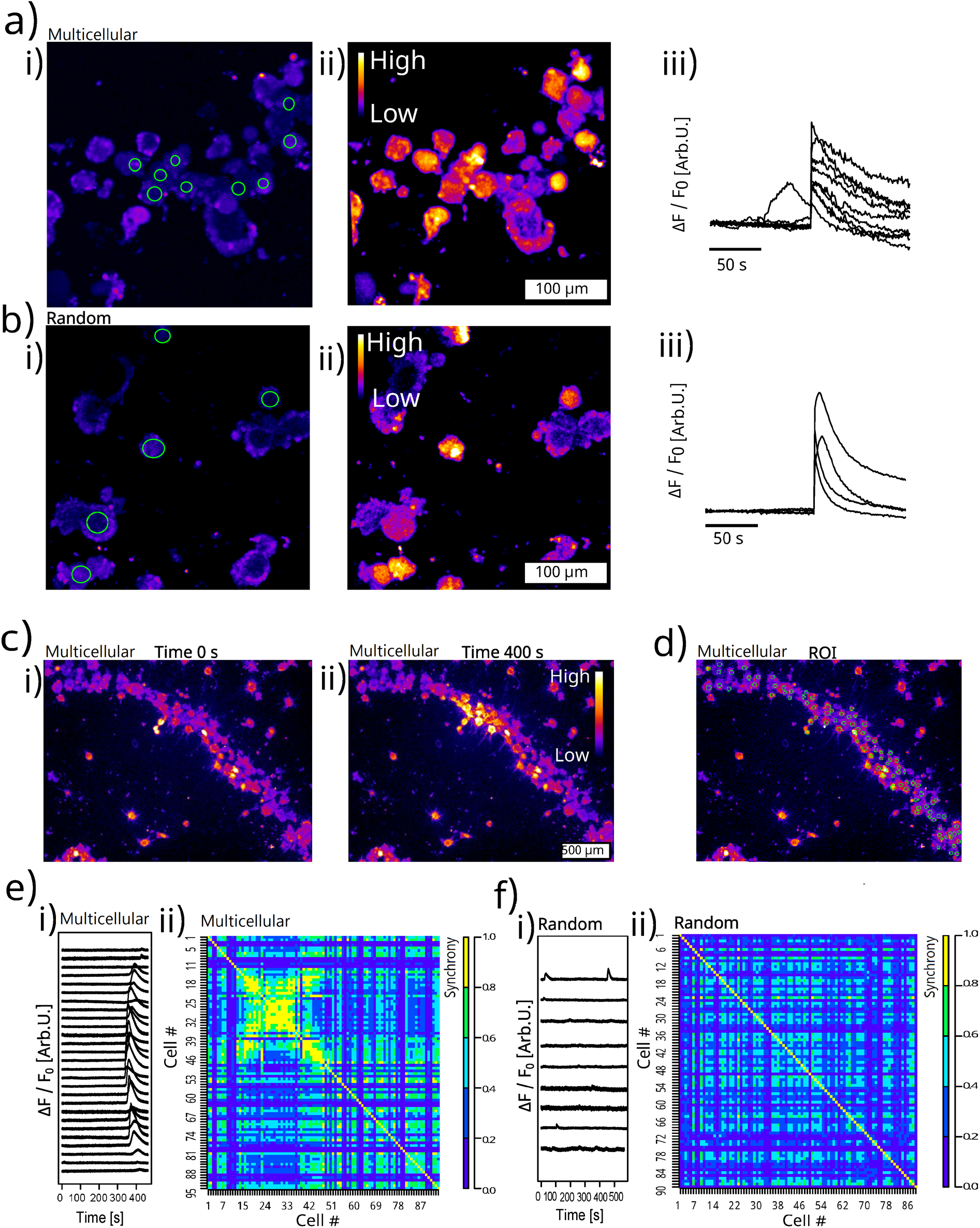
Calcium signal evaluation. **a-b)** Calcium imaging shows that neurons are functional when depolarized by 50 mM potassium chloride at day 2. **c)** Neurons in the multicellular system evolved a functional crosstalk after 4 days of culture. The calcium signal is transmitted from one cell to its neighbors only in the multicellular system. The color scale of a-c) are ΔF / F_0_. **d)** The region of interest (ROI) of the tissue units. **e-f)** The multicellular system (e) displays higher synchrony of calcium signal in the tissue units than in its random counterpart (f). (i) Normalized calcium fluorescent curves. Only tissue units with spontaneous calcium signals are presented. (ii) The synchrony matrix.

Despite the activation of individual neuron, the cellular function at the population level is largely unveiled. *In vivo* DRG exhibits functional crosstalk within the network of neurons and SGCs. The calcium signals can transmit from cell to cell involving both SCGs and neurons.^[17a]^ Our multicellular system of DRG recapitulates this *in vivo* functional crosstalk. The calcium signals travel over a distance of 200-500 μm, crossing 5-20 tissue units (Figure 3c). When the active fluorescent traces are extracted from regions of interest (ROIs) of the DRG tissue units, the spikes of calcium signals are exclusively synchronized in the multicellular system (Figure 3e-f (i)). These synchronized calcium signals are observed only in the multicellular system, but never in experiments of random culture (n=4 experiments per group). We calculate the synchrony of tissue units which is defined as cross-correlation coefficient at zero lag time.^[27]^ The synchrony matrix shows a high degree of synchronization in a subpopulation of closely packed tissue units (note the yellow region in the matrix of Figure 3f (ii)), while an evident lower synchrony is found in the random culture (Figure 3e (ii)). The results demonstrate that the multicellular morphology can be engineered to reproduce multicellular crosstalk and to develop populational-level functions.

### 2.3. Morphology of the DRG multicellular system and its self-organization

Neurons are densely packed in native DRG tissue (Figure 4a). This close cell-to-cell proximity is successfully recapitulated in the hydrodynamic force-assembled multicellular system (Figure 4b). The neuron sizes in the assembled multicellular system approximate those of bovine DRG native tissue (Figure 4c-d).^[9]^

**Figure 4.**
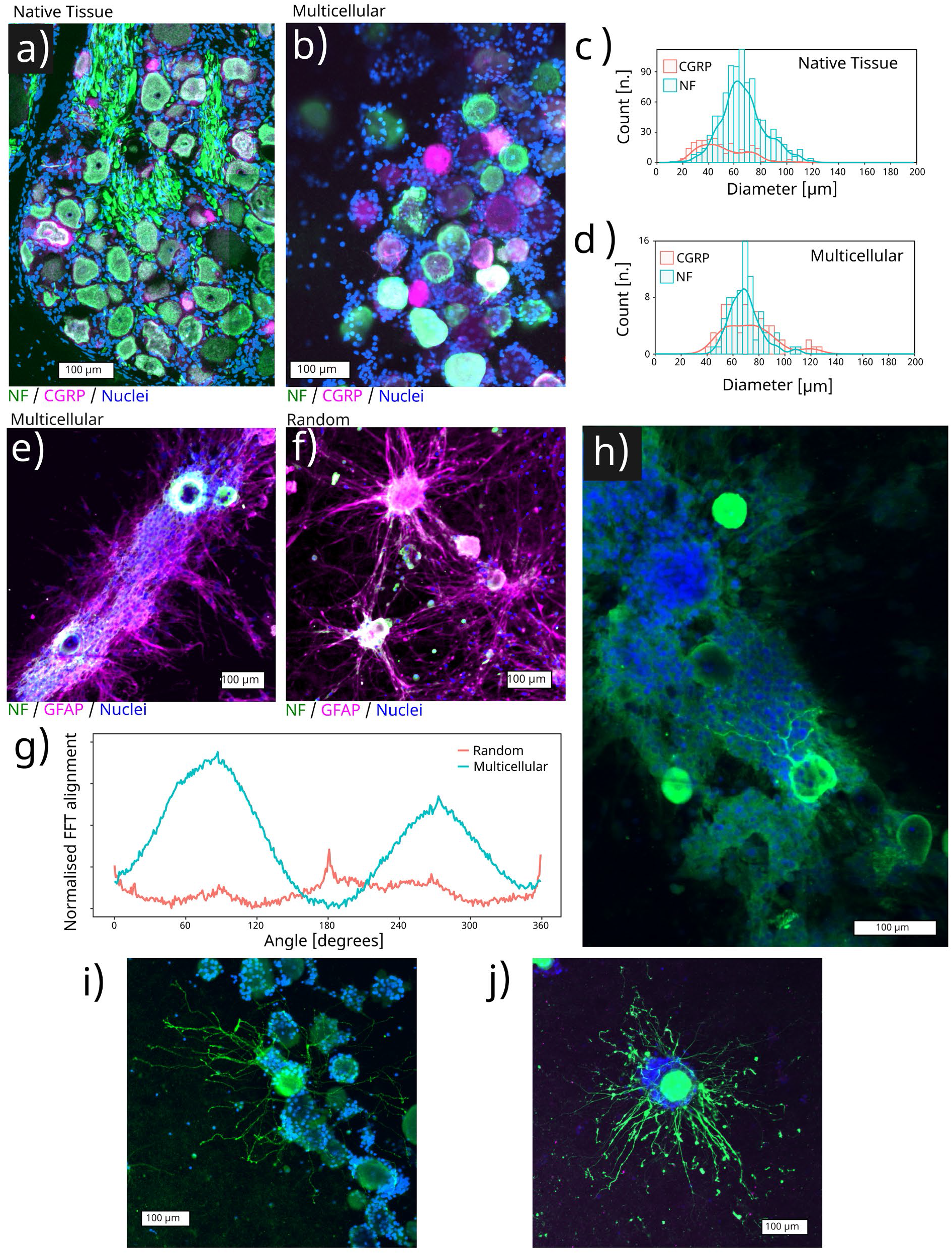
Morphology of the assembled multicellular system. Immunofluorescent staining of the native tissue **(a)** and of the assembled multicellular system **(b)** which closely recapitulate the tight cell-to-cell contact of the native DRG. Soma diameters were comparable between the multicellular culture **(c)** and the native bovine DRG tissue **(d)**. The multicellular assembly presented anisotropic cells’ self-organization **(e)**, while in the random culture the self-organization of satellite glial cells appeared limited at day 8 **(f)**, as evidenced by the quantification done via the 2D-FFT alignment **(g)**. The self-organized structure provides a guidance for axonal growth **(h)**. Axons sprouting in collagen matrix in both the multicellular system **(i)** and the random culture **(j)** after 2 days of culture.

Native DRG tissue is also known to have oriented cellular anisotropy. Figure 4a shows that the SGC nuclei are oriented in the same direction of axon fibers. Moreover, we show that the self-organization capability is not limited to stem cells. Mature DRG cells also exhibit the ability to self-organize. At day 8, SGCs in the multicellular system orient themselves along a precise direction connecting each other in an organized structure (Figure 4e). Importantly, in the random culture SGCs display only a small trend to self-organize. Merely few SGCs tend to extend and connect isolated DRG tissue units (Figure 4f). This is further confirmed by the 2D-FFT analysis. The orientation distribution plot shows a homogeneous SGC alignment in the multicellular system, while the orientation distribution of non-assembled control is mostly random (Figure 4g). Such anisotropic SGCs organization seems to guide and support axonal growth, similarly to what observed in native DRG tissue (Figure 4a and h).

Next, to study the sole effect of AF explant on nerve ingrowth and avoid the interference of self-organizing SGCs, we use the day-2 culture to evaluate the ingrowth-associated axonal sprouting. At day 2, the SGCs are still symmetrically enveloping the neuronal soma without any oriented spreading. Axons extend in the collagen matrix without the influence of SGCs (Figure 4i-j).

### 2.4. The neurotrophic (growth-promoting) effect of cytokine-primed AF depends on the AF-neuron distance

We prove the neurotrophic effect of the cytokine-primed AF on the CGRP(+) neurons when the AF-neuron distance is smaller than 1300 μm (AF-close neurons). For AF-close and CGRP(+) neurons, the average axonal length of cytokine-primed AF group is ∼25 μm longer in the random culture (Figure 5a (i-ii)), and ∼27 μm longer in the multicellular system (Figure 5b (i-ii)) compared to the non-cytokine control.

**Figure 5.**
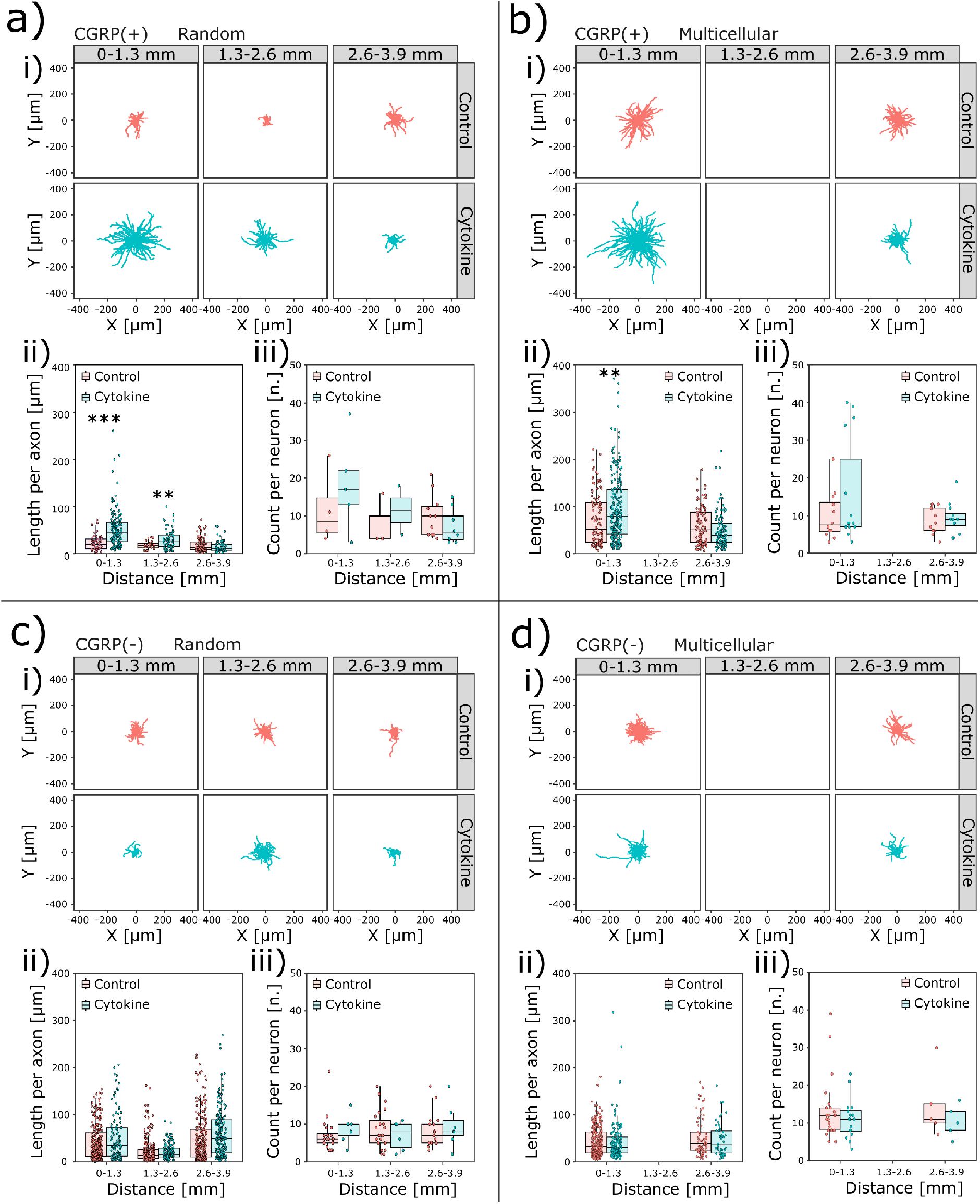
Effect of cytokine-primed AF tissue on axonal length and count depends on the AF-neuron distance. Axonal morphologies are shown as a function of the distance from the AF tissue in a-d (i). CGRP(+) axons in both random culture **(a)** and multicellular system **(b)** show a distance-dependent increased length (ii) and count (iii) when exposed to a cytokine treated AF tissue. CGRP(-) and NF(+) axons do not show any distance-dependent difference in length (i) or count (ii) in neither random culture **(c)** nor multicellular system **(d)**. **: p<0.01, ***: p<0.001.

We find this neurotrophic effect of cytokine-primed AF depends on the AF-neuron distance. A closer AF-neuron distance is associated with a longer axonal extension, thus displaying a negative correlation between the AF-neuron distance and the length of CGRP(+) axons. The correlation coefficient (ρ) between the AF-neuron distance and CGRP(+) axon length is −0.55 (p<0.001) in the cytokine-primed AF group (Figure 5a (ii green)). The close AF-neuron distance is also correlated with a higher count of CGRP(+) axons (axonal branch number) exhibiting aρvalue of −0.52 (p=0.03) (Figure 5a (iii green)). The AF-far neurons are less influenced by the cytokine-primed AF explant and can be used as a control. When AF is not primed with cytokines, the axonal length and count of AF-close neurons are not different from those of AF-far neurons. Theρvalues between AF-neuron distance with CGRP(+) axonal length and with CGRP(+) axonal count are −0.16 (p=0.03) and 0.08 (p=0.75), respectively (Figure 5a-b (i-iii red)). Thus, there is no detectable neurotrophic effect in the non-cytokine control.

These results point to a need to control the AF-neuron distance. The hydrodynamic forces assemble a larger number of neurons to be AF-close and potentially increases the sample size of neurons for analysis. This is of practical importance. In the random culture, many experiments without any neurons close to the AF tissue have to be excluded. After performing the random culture 96 times using DRG from 12 cattle, only 9 CGRP(+) neurons with 186 axons were obtained in the AF-close region. Instead, the assembled culture achieved a sample size of CGRP(+) neuron more than 2-fold higher than the random culture in the AF-close region with only 8 experiments using DRG from 4 cattle. In the assembled multicellular system that is AF-close, 27 CGRP(+) neurons with 355 axons were available for analysis (Figure 5a-d (ii) number of data points).

For CGRP(-) neurons, the AF explant is not showing significant neurotrophic effect. The CGRP(-) axon length and count are not different comparing AF-close and AF-far neurons and are not influenced by the cytokine-priming of AF explant (Figure 5c-d (i-iii)). This agrees with the *in vivo* findings where axons growing into pain-generating IVD are usually labelled by CGRP(+) or SP(+).^[7b, 10b]^ Those CGRP(-) / SP(-) and neurofilament (NF)(+) neurons are regarded as low-threshold mechanoreceptors and proprioceptors that sense light touch and body position.^[7d]^ CGRP and NF are specific markers of peptidergic nociceptors and non-nociceptors, respectively. These 2 markers are shared between species.^[7d]^ Based on the results of our model, the neurotrophic effect of cytokine-primed AF is nociceptor specific.

### 2.5. The neurotropic (guidance) effect of cytokine-primed AF

To evaluate how far the CGRP(+) axon can grow towards AF, we projected the axon trajectory to the axis connecting AF and neuron cell body to measure the projection length. The maximum projection length of axons per neuron was evaluated (Figure 6a (i)). A larger maximum projection length indicates that the axons extend more towards the AF. Results imply that the AF’s influence on maximum projection length also depends on the AF-neuron distance. In the cytokine-primed AF group, the ρvalues are −0.59 (p=0.012) and −0.30 (p=0.12) for random culture and multicellular system, respectively (Figure 6a (ii-iii green)). However, in the non-cytokine control, such correlation is not observed. The ρvalues are 0.15 and −0.02 for random culture and multicellular system, respectively (Figure 6a (ii-iii red)). These results indicate that the cytokine-primed AF may have a neurotropic effect on CGRP(+) and AF-close axons.

**Figure 6.**
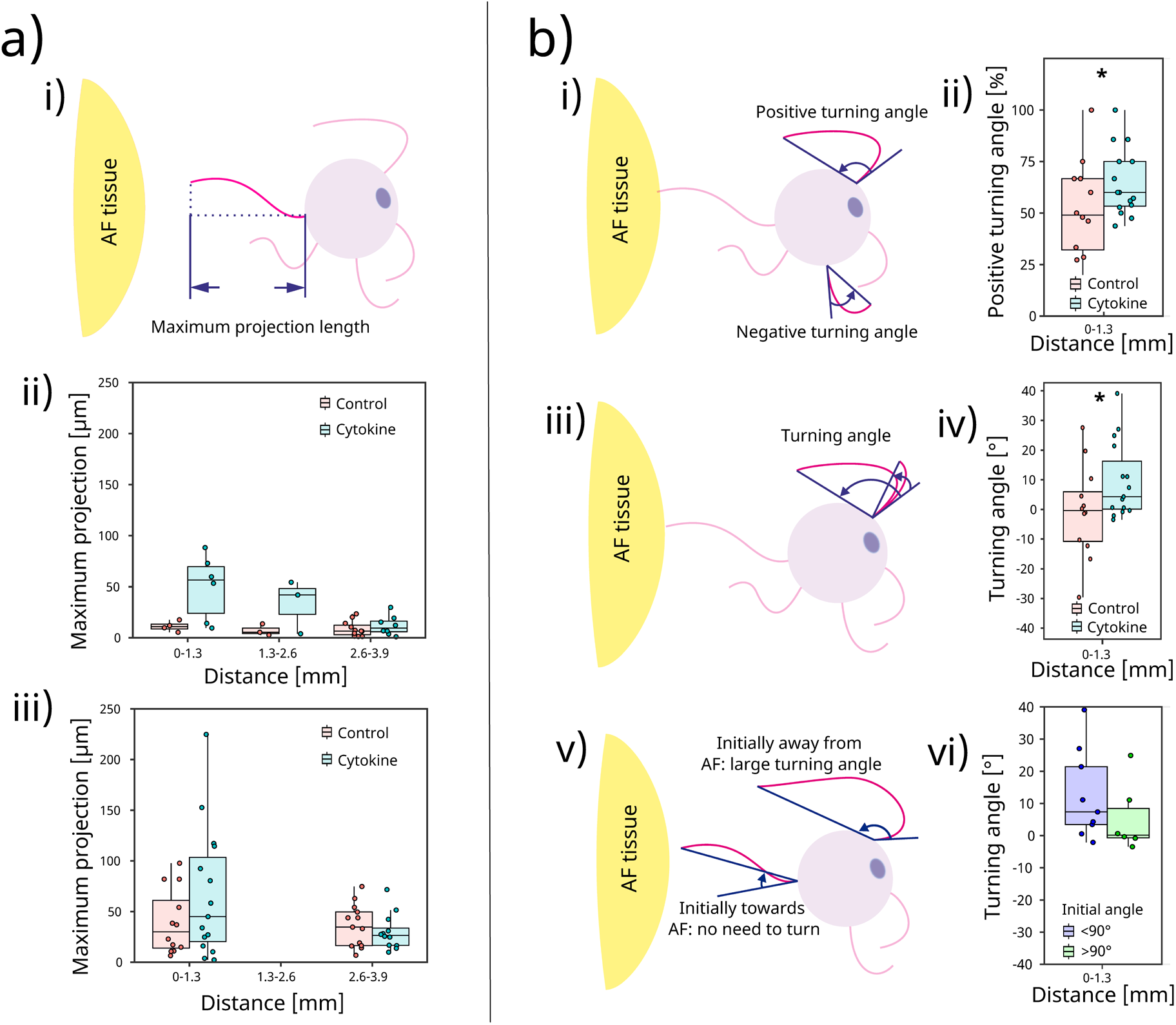
Effect of the cytokine treated AF tissue on axonal guidance. **a)** Schematic depiction illustrating the ‘maximum projection length’. (i). CGRP (+) axons showed a distance-dependent ‘maximum projection length’ in both random culture (ii) and multicellular system (iii). **b)** Turning angle of axons. (i) ‘Positive’ and ‘negative’ turning angles. (ii) A higher proportion of axons per neuron turned towards the cytokine-treated AF tissue, but not towards the non-cytokine-treated control. iii) Illustration showing larger turning angle correspond to a larger AF guidance. iv) Cytokine-treated AF tissue induces a larger turning angle than the control. As illustrated in (v) and quantitatively shown in (vi), the guidance on axons results in a larger turning angle only when axons initially sprout in the opposite direction in respect to AF (initial angle > 90°). *: p<0.05.

The developing nervous system is far more plastic (capability to change) than the adult, so embryonic and neonatal neuron culture are more commonly used to investigate how axons sense the environmental cues and decide the growing direction.^[28]^ Here, we evaluate the neurotropic effect of AF on mature neurons which has seldomly been characterized before. Since the random culture did not have enough neurons (9 AF-close neurons), this analysis was only possible for the multicellular system (27 AF-close neurons). The turning angle is a common indicator of axonal guidance.^[28d, 28e]^ It is defined by the difference between the initial growth angle and the ending growth angle (Figure 6b (i, iii, v)). An AF’s guidance on attracting axon will result in a positive turning angle, while an axon turning away from the AF explant is represented by a negative turning angle (Figure 6b (i)). In the non-cytokine control, the proportion of positive turning axons per neuron averages at 50% suggesting equal numbers of axons turning toward and away from AF. Thus, there is no guidance from the AF tissue (Figure 6b (ii red)). In contrast, in the cytokine-primed AF group, 60% of CGRP(+) axons per neuron have a positive turning angle. Notably, in the cytokine-primed AF group, 12 out of 14 CGRP(+) neurons (85.7%) have majority of their axons turning with a positive angle (Figure 6b (ii green)).

The turning angle value is averaged to be −0.4° for non-cytokine control but is 4.3° for the cytokine-primed AF group (Figure 6b (iv)). The axonal guidance is represented by a larger turning angle only when the axon is initially pointing away from the AF (Figure 6b (iii)). Nevertheless, when the axons initiated their growth already towards the AF, the AF’s guidance does not need a turning, but instead to guide the axons to persist the initially ‘correct’ direction (Figure 6b (v)). Considering this, we divide the data points of cytokine-primed AF group into initial angle greater than 90° and smaller than 90°. We do find a larger turning angle when the axons are initially pointing away from the AF (initial angle < 90°) compared to axons initially already pointing towards the AF (initial angle > 90°) (Figure 6b (vi)). Taking together, the cytokine-primed AF shows an ingrowth-related axonal guidance.

## 3. Discussion

Chronic low back pain (LBP) is affecting more than 500 million people and is the leading cause of disability worldwide.^[29]^ Although most structures in the lumber region can contribute to LBP, the dysfunctional intervertebral disc (IVD) is recognized as the source of pain in 40% of LBP patients.^[30]^ The associated sensory nerve ingrowth in IVD may contribute to chronic pain.^[7b]^ Our model reveals that the pro-inflammatory cytokine-primed AF tissue has growth promoting (neurotrophic) and chemotactic (neurotropic) effects in large animal mature neuron system. The model may be useful to study the mechanism of IVD nerve ingrowth for a better understanding of low back pain.

To model inter-tissue/organ communication, recapitulating the *in vivo* anatomical proximity is essential. In the case of IVD, the nociceptor axons are closely attaching to the outside collagen matrix of AF *in vivo*.^[6]^ Indeed, we found that only when this AF-nerve apposition is recapitulated, nociceptor axons can be influenced by cytokine-primed AF. Also, our model is based on type Ⅰ collagen matrix which mimicry the *in vivo* condition.^[16, 31]^ Remarkably, the adaptability of our experimental set-up can be explored to model other tissue nerve ingrowth, provided that the corresponding morphologically and physiologically relevant environments are recapitulated.

Most *in vitro* study of DRG neurons are based on primary monolayer culture and cell line culture. Neurons are cultured on stiff plastic and glass surface and are subject to a different mechanical environment comparing to the *in vivo* situation.^[32]^ Culturing DRG neurons in extracellular matrix-based hydrogel is more physiological,^[32a]^ but cells are still isolated, losing the multicellular architecture of native tissue. Although culturing tissue explant *ex vivo* preserves the tissue architecture,^[31]^ explant culture is only possible for the study of rodent embryonic and neo-natal DRG.^[33]^ The culture of large animal or human DRGs in the form of tissue explant is challenging and has not yet been reported. Large animal or human DRGs are much larger than rodent DRG, and the extracellular matrix is denser preventing nutrients ^[9]^ and oxygen diffusion.^[34]^

Our approach can overcome these limitations. Large animal DRGs are first disintegrated into micro-scale tissue units, and then re-assembled using mild acoustic hydrodynamic forces. In this way, DRG are maintained in a multicellular system with a controlled scale to reduce the risk of necrotic core. In our DRG multicellular system, the dimension of the assembled system can be controlled in a width of 300-500 μm (Figure 2 and S2) and a thickness of 100-150 μm. The physiological culture condition of our multicellular system is evidenced by the high cell viability.

Former study showed that acoustic assembled embryonic mouse cortical brain cells exhibit inter-neuron synchronization.^[22b]^ Little is known whether this shape-to-function relationship also exist in adult mature neurons. Our model shows a cell-building morphology cue in inter-neuronal crosstalk in the case of mature neurons. The inter-neuronal crosstalk in DRG has been increasingly recognized as an essential mechanism of chronic pain^[17a]^. This knowledge may contribute to the improvement of the general DRG culture protocol, where the neural cells can be investigated in a populational level considering cell-to-cell communication and network.

Over decades, embryonic and neo-natal DRG from chicken, mice and rat has greatly contributed to the understanding of sensory neural biology. However, the increasing call for translation from benchside to bedside requires models with higher clinical relevance. We must admit that these lower vertebrates (*e*.*g*., rodents) are biologically different from human^[35]^. Also, the developing (embryonic and neonatal) nervous system is different from their adult and mature counterpart.^[36]^ Likewise, the maturity of neuron-like cells derived from human induced pluripotent stem cells is not yet fully validated, and little is known about their similarities to mature native DRG neurons.^[14b]^ Experiments using adult human DRG are constrained by the limited availability and ethical considerations. Recently, researchers are beginning to focus on adult large animal DRG to perform transcriptome and to map genomic loci associated with human pain disease.^[7d]^ Here, large animal DRG and AF are co-cultured to model pathophysiology associated with human disease. Our adult large animal tissues are obtained from the abattoir, with cheap, unlimited resource and no ethical concern. This large animal model may leverage the strength of bio-engineering technologies in studying human relevant disease.

## 5. Conclusion

We reported the first *in vitro* model of AF nerve ingrowth using large adult animal tissue units. Mild hydrodynamic forces are used to spatially assemble DRG micro-scale tissue units in a defined spatial organization. We proved the importance of recapitulating the *in vivo* morphology for developing physiological functions. By assembling DRG tissue units into a densely packed multicellular system, we achieved native-like structural organization that led to multi-level functional crosstalk (inter-cellular and inter-tissues). We demonstrated that the influence of cytokine-primed AF explant on the extension and guidance of CGRP(+) axons depends largely on the AF-neuron distance. This underpins the role of reproducing inter-tissue/organ anatomical proximity when investigating their communications. This work proposes a new approach to biofabricate multi-tissue/organ models for unraveling clinically relevant pathophysiology as well as discovering novel treatments.

## 4. Experimental Section

### AF tissue dissection, culture, and pro-inflammatory cytokine priming

Bovine tail IVDs were obtained from the local abattoir. Donor information (sex, age and weight) is in Table S1. The IVDs were transected at cartilage growth plate, cleaned by a jet lavage and rinsed in phosphate buffered saline (PBS) containing 10% penicillin-streptomycin (PS, 15140-122, Gibco, UK). After removing the IVD endplates, the outer AF tissue was cut into small pieces with thickness ranging from 1 to 50 mm for AF viability essay (Figure S1) and 1.5×1.5×1.5 mm cube for the co-culture with DRG tissue units. The AF tissues were maintained in 4.5-g/L glucose Dulbecco’s Modified Eagle Medium (DMEM, 52100-021, Gibco, Paisley, UK), 10% fetal calf serum (FCS, 35-010-CV, Corning, CA, US), 1% PS and 0.11-g/L sodium pyruvate (P5280; Sigma-Aldrich, JP) until use. The viability of the cultured AF tissue was evaluated by live dead staining. To allow dye penetration, the AF explants were pre-processed by 1 mg/mL collagenase P (Roche, Mannheim, DE) for 1 h at 37 °C. The dyes used for live dead staining were Calcein-AM (1 µM, 17783, Sigma) and Ethidium homodimer-1 (1 µM, 46043, Sigma). The AF explants were incubated in the day for 30 min at 37 °C. The imaging was performed using LSM 800 confocal (Zeiss, DE).

### Bovine DRG micro-tissue unit preparation

Adult bovine DRGs were obtained from cervical spines at the local abattoir. Donor information is in Table S1. 10-12 DRGs per donor were disintegrated into micro tissue units using 6 mL collagenase P (4 mg/mL, Roche, Mannheim, DE) in a 15 mL Falcon tube on an orbital shaker for 2 h at 37 °C. A manual shaking of the digesting tissue was performed every 15 min. The digested compound went through a 100 μm cell strainer (Falcon, Corning, US) and centrifuged through 5 mL of PBS containing 15% BSA (centrifugation speed at a speed 600 g and duration for 7 min) in a 50 mL Falcon tube. The pellet was re-suspended in 500 μL DMEM/F12 (50% v/v, DMEM from Gibco, 52100-021, UK and F-12 Ham from Sigma, N6760, UK) supplemented with 10% FCS, 20 mM HEPES (15630122, Thermo Fisher, US) and 1% ITS (354352, Sigma-Aldrich, MO, USA). 50 μL of the re-suspended solution was stained by equal amount of 0.4% trypan blue (T8154, Sigma) and counted in a hematocytometer.

### Collagen matrix preparation

Collagen solution containing DRG tissue units was prepared by sequential mixing of the following components: 10% of 10× concentrated DMEM/F12, 10% (v/v) FCS, 1% (v/v) ITS, 20 mM HEPES, 62% (v/v) deionized water, 0.5 mg/mL Collagen I (stock at 10 mg/mL, cat.no. 354249, Corning, MA, US), and 10% (v/v) DRG tissue unit solution (1.2 × 10^5^ units/mL). For each 100 μL stock collagen solution (10 mg/mL), 6 μL 7.5% sodium bicarbonate was added to adjust the pH. This preparation was on ice and the sound-assembling was performed immediately following the solution preparation.

### Hydrodynamic assembly of DRG micro-tissue units around AF explant

The AF explants were trimmed to cubes sized at 1.5×1.5×1.5 mm and were embedded in 360 μL of 2 mg/mL collagen in a culture frame. After 12h, assembly was performed on top of the AF-embedding collagen in a shallow cylindrical space with a diameter of 25 mm and height of 2 mm. The volume of the DRG containing collagen solution was 575 μL. The assembly protocol is based on former publication^[22a]^. Briefly, the excitation frequency was 60 Hz and amplitude 0.5 g. The duration of assembly was 2 min. Once collagen formed a hydrogel, the frames were moved to a 37 °C incubator supplied with 90% humidity and 5% CO_2_.

### Immunofluorescence

For CGRP and NF labelling, the primary antibodies included rabbit anti-CGRP antibody (1:500 dilution volume, cat.no. 24112, Immunostar, Sodiag Avegno, WI, US) and mouse anti-neurofilament 200 antibodies (NF200, 1:50 dilution, cat.no. OMA1-06117, Thermo scientific, NL). The primary antibody incubation was at 4 °C for 24 h. After washes, this was followed by 2 h of secondary antibody incubation. The secondary antibodies included goat anti-mouse AlexaFluor 488 conjugated antibody (1:100, A-11029, Thermo Fisher, OR, US) and donkey anti-rabbit AlexaFluor 680 (1:100, A32802, Thermo Fisher, OR, US). Nuclei were stained using Hoechst (2 μM, Sigma, cat. n. 14530, US). The gel volume was included for the calculation of antibody dilution. LSM-800 confocal was used for imaging. Z-stack was performed to ensure that the whole length of axons was covered in the images. For GFAP labelling of SGCs combined with neuron labelling (NF and TUBB3), the primary antibodies included rabbit anti-GFAP polyclonal antibody (1:200, Dako, CA, USA) and mouse anti-TUBB3 (1:50, MAB1637, Sigma, CA, USA). The staining and imaging method was the same as the labelling of CGRP and NF. The anisotropy of SCGs was characterized by the formerly reported 2D-Fast Fourier Transform (FFT) method.^[37]^ Briefly, the intensity profiles were obtained summing all the pixel fluorescent intensity (I) along a column of the image and normalizing the intensity profile to 0.

### Calcium imaging

The calcium imaging protocol has been described formerly.^[38]^ Briefly, calcium imaging was performed using 5 μM Fluo-4-AM (ThermoFisher, F14201, US) in Krebs-Ringer’s solution containing NaCl 119 mM, KCl 2.5 mM, NaH_2_PO4 1.0 mM, CaCl_2_ 2.5 mM, MgCl_2_ 1.3 mM, HEPES 20 mM and D-glucose 11.0 mM. Time-lapse images were acquired using a LSM800 microscope with wide field epi-fluorescent camera at 5× magnification and 5×5 binning. The image acquisition rate was 30Hz. Calcium imaging videos were further analyzed with ImageJ Fiji (1.53q NIH, US) based on java 1.8.0_322 (64-bit).

The regions of interests (ROIs) were manually selected, and the fluorescence intensity was extracted from each ROI. The fluorescence intensity data was then imported into ‘R studio’ for further analysis. The baseline fluorescence signal (F_0_) was defined as the average fluorescent intensity of the first 1000 time points. The real-time fluorescent intensity was normalized: ΔF / F_0_ = (F-F_0_) / F_0_. The peaks of ΔF / F_0_ were detected as local maximum in each ROI. A spontaneous calcium signal was defined as a peak height 4 × standard deviation (S.D.) higher than the baseline level. The synchrony between ROIs was evaluated following formerly reported methodology.^[27]^

### Data analysis

The image segmentation and measurement were performed using imageJ Fiji (1.53q NIH, US) based on java 1.8.0_322 (64-bit). Simple neurite tracer plugin (SNT)^[39]^ was used to segment axons in z-projected 3D z-stacked images. The segmented axons were saved in ‘swc’ format and imported into ‘R studio’ (1.1.383) based on ‘R’ (3.6.2) using the ‘nat’ package. Data arrangement and statistic were performed using ‘tidyverse’ package within R. Visualization of the data was performed using ‘ggplot2’ package.

## Author contributions

J. Ma: Methodology, Investigation, Data curation, Formal analysis, Writing – original draft, review & editing; J. Eglauf: Investigation; R. Tognato: Methodology, Data curation, Visualization, Formal analysis, Writing – review & editing; S. Grad: Writing – review & editing; M. Alini, T. Serra: Conceptualization, Funding acquisition, Supervision, Writing – review & editing.

## Acknowledgements

The presented work is funded by AO Foundation (Davos, Switzerland).

