## Supporting information for "Engineering sensory ganglion multicellular system to model tissue nerve ingrowth"

J. Eglauf

ETH Zürich, Rämistrasse 101, 8092, Zürich, Switzerland

T. Serra

MERLN Institute for Technology-Inspired Regenerative Medicine, Maastricht University

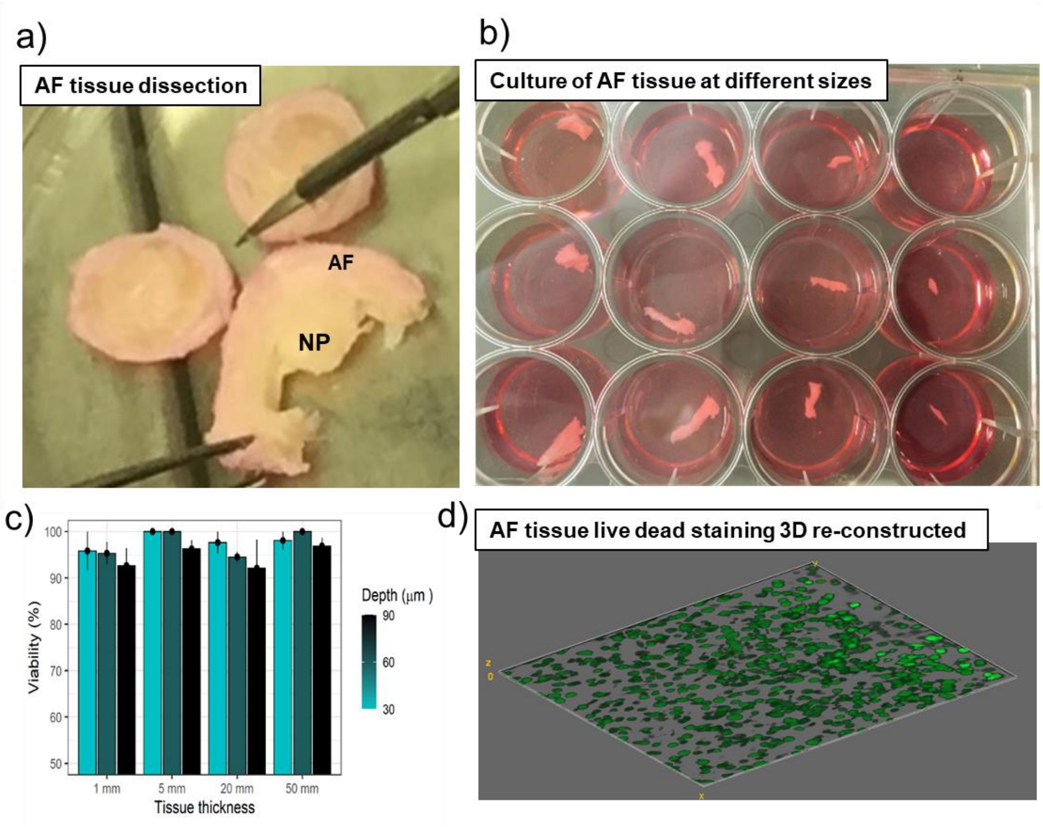


**Figure S1.** The culture of AF tissue explant. **a)** Dissection of AF tissue. **b)** Outer AF tissues are trimmed into different sizes for culture. **c)** The live dead staining of AF tissue after 9 days of culture shows a viability higher than 90% and this is not influenced by the trimming size of the explant. **d)** 3D reconstruction of confocal z-stack images of the live-dead-staining AF cells to 90 µm depth from the surface of AF explant.

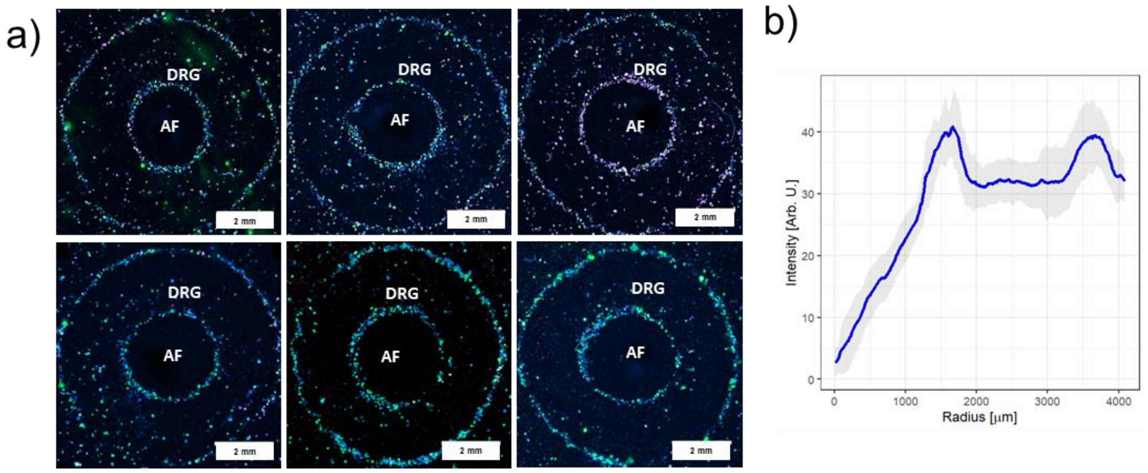


**Figure S2.** Geometry of the hydrodynamic patterning. **a)** Six experiments showing the consistency of the patterned DRG multicellular system. These are tiled immunofluorescent images labelling NF200 (green), CGRP (purple) and nuclei (blue). **b)** Radial profile of the patterned cellular geometry evaluated as in Di Marzio *et al.* [1] The 2 peaks represent the 2 rings, and the width of the peaks indicates the ring thickness. The x-axis location of the peaks shows diameter of the rings. Grey stratum represents standard deviation among different experiments.

**Table S1.** Donor information of bovine annulus fibrosus tissues (AF) and bovine dorsal root ganglion tissues (DRG)

| Donor ID | Sex | Age (month) | Weight (kg) | Tissue Type |
|---|---|---|---|---|
| #1 Random[a)] | Female | 12 | 200 | AF |
| #2 Random | Male | 10 | 190 | AF |
| #3 Random | Male | 10 | 190 | AF |
| #4 Random | Male | 10 | 190 | AF |
| #5 Random | Male | 6 | 148 | AF |
| #6 Random | Male | 12 | 200 | AF |
| #7 Random | Male | 12 | 192 | AF |
| #8 Random | Male | 5 | 132 | AF |
| #9 Random | Female | 6 | 195 | AF |
| #10 Random | Male | 4 | 130 | AF |
| #11 Random | Male | 4 | 122 | AF |
| #12 Random | Male | 4 | 122 | AF |
| #13 Random | Male | 11 | 240 | DRG |
| #14 Random | Male | 10 | 210 | DRG |
| #15 Random | Male | 12 | 204 | DRG |
| #16 Random | Male | 12 | 204 | DRG |
| #17 Multicellular[b)] | Male | 13 | 230 | AF |
| #18 Multicellular | Male | 12 | 235 | AF |
| #19 Multicellular | Female | 13 | 244 | AF |
| #20 Multicellular | Female | 8 | 180 | AF |
| #21 Multicellular | Female | 6 | 132 | DRG |
| #22 Multicellular | Female | 10 | 169.8 | DRG |

[a)] Random: the DRG tissue units seeded randomly around AF without assembling; [b)] sound-assembled DRG into multicellular system around AF.
